# Urine proteome changes in rats with approximately ten tumor cells subcutaneous inoculation

**DOI:** 10.1101/604520

**Authors:** Jing Wei, Wenshu Meng, Youhe Gao

## Abstract

Biomarkers are changes associated with the disease. Without homeostatic control, urine accumulates very early changes and is an ideal biomarker source. Usually, we performed urinary biomarker studies involving at least thousands of tumor cells. But no tumor starts from a thousand tumor cells. Can we observe any urine proteome changes in rats with approximately ten tumor cells subcutaneous inoculation? Here, we serially diluted Walker-256 carcinosarcoma cells to a concentration of 10^2^/mL and subcutaneously inoculated 0.1 mL of these cells into nine rats. Urine proteomes on days 0, 13 and 21 were profiled by LC-MS/MS analysis and studied with unsupervised clustering analysis. Samples at three time points were almost clustered together, indicating a good consistency in these nine rats. Differential proteins on days 13 and 21 were mainly associated with cell adhesion, autophagic cell death, changes in extracellular matrix organization, angiogenesis, and the pentose phosphate pathway. All of these enriched functional processes were reported to contribute to tumor progression and could not be enriched through random allocation analysis. Our results indicated that 1) the urine proteome reflects changes associated with cancer even with approximately ten tumor cells in the body and that 2) the urine proteome reflects pathophysiological changes in the body with extremely high sensitivity and provides potential for a very early screening process of clinical patients.

## Introduction

Urine is an ideal biomarker resource. Without homeostatic mechanisms, urine can accumulate all changes from the whole body; therefore, urine may have the potential to detect early and small changes in the body^[1]^. Urine is easily affected by various physiological factors, such as sex, age and diet^[2]^. In patients, the urine proteome is easily influenced by some medications because therapeutic measures for patients are inevitable. Therefore, our laboratory proposed a strategy for urinary biomarker studies. First, we used animal models to find early biomarkers of related diseases. Then, we verified candidate biomarkers in clinical urine samples^[3]^. The use of animal models minimizes external influence factors, and differential urinary proteins found in animal models are more likely to be directly associated with related diseases. According to this strategy, our laboratory has applied different types of animal models, such as subcutaneous tumor-bearing model^[4]^, pulmonary fibrosis^[5]^, glioma^[6]^, liver fibrosis^[7]^, Alzheimer’s disease^[8]^, chronic pancreatitis^[9]^ and myocarditis^[10]^, to search for early biomarkers before the appearance of pathology changes and clinical manifestations.

Urine can reflect not only early changes in related diseases but also more changes than that in blood. When introduced the interference into the blood by using two anticoagulants, more proteins and more information on the changes in proteins were detected in urine than in plasma^[11]^. In our previous tumor-associated studies, such as in the subcutaneous tumor-bearing model^[4]^ and glioma rat model^[6]^, millions of tumor cells were injected into the animal models. Since urine is a more sensitive biomarker resource than blood, we explored the sensitivity limit of urine. If there were only limited tumor cells in the body, for example, approximately ten tumor cells, can the urine proteome reflect the changes?

In this study, we subcutaneously injected approximately ten Walker-256 carcinosarcoma cells into nine rats. Urine samples were collected on days 0, 13, and 21. Urine proteins were analyzed by liquid chromatography-tandem mass spectrometry (LC-MS/MS). Differential proteins on days 13 and 21 were analyzed by functional enrichment analysis to find associations with tumor progression. This research aimed to determine whether the urine proteome could reflect changes associated with these ten tumor cells. The technical flowchart is presented in Figure 1.

**Figure 1.**
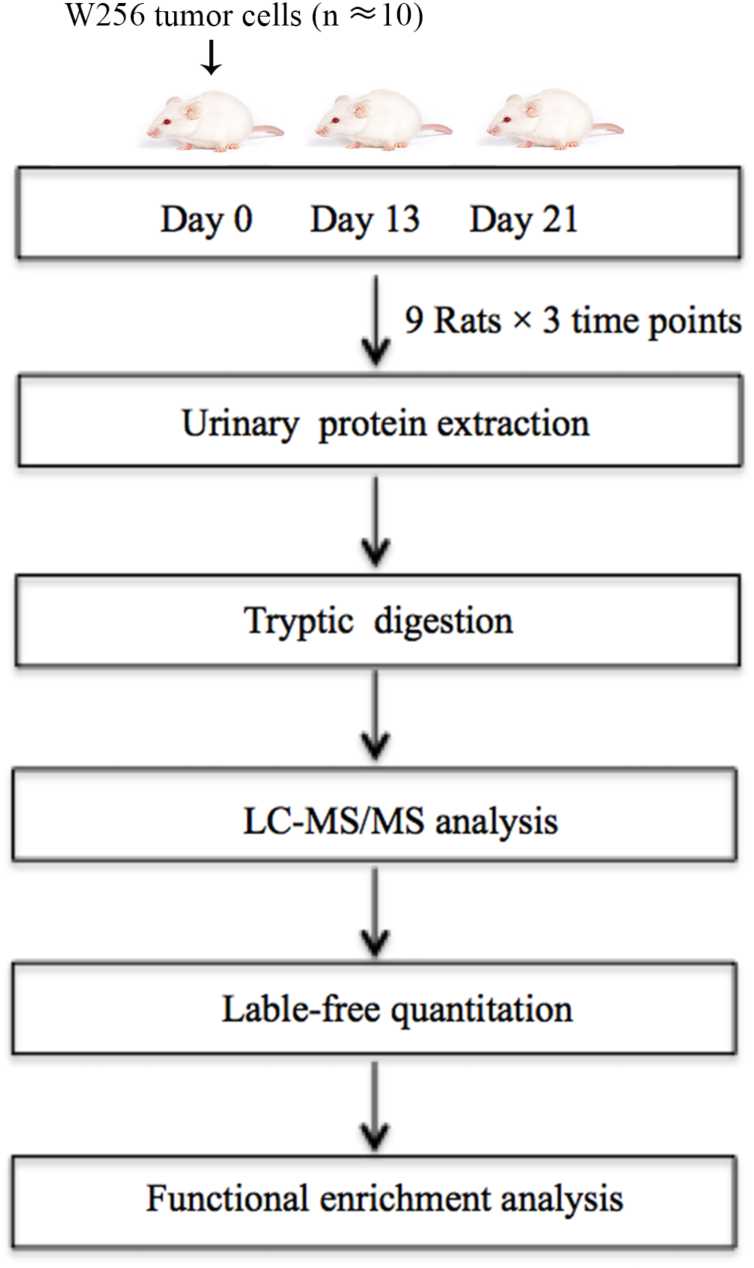
Workflow of protein identification in ten tumor cells subcutaneously inoculated in rats. Urine was collected on days 0, 13 and 21 after inoculating tumor cells. Urinary proteins were extracted, digested, and identified by liquid chromatography coupled with tandem mass spectrometry (LC-MS/MS) identification. Functional enrichment analysis of differential proteins was performed by DAVID and IPA.

## Materials and methods

### Animal treatment

Male Wistar rats (n=12, 150 ± 20 g) were purchased from Beijing Vital River Laboratory Animal Technology Co., Ltd. Animals were maintained with a standard laboratory diet under controlled indoor temperature (21±2°C), humidity (65–70%) and 12 h/12 h light–dark cycle conditions. The experiment was approved by Peking Union Medical College (Approval ID: ACUC-A02-2014-008).

Walker-256 (W256) carcinosarcoma cells were purchased from the Cell Culture Center of Chinese Academy of Medical Sciences (Beijing, China). W256 tumor cells were intraperitoneally inoculated into Wistar rats. The W256 ascites tumor cells were harvested from the peritoneal cavity after seven days. After two cell passages, the W256 ascites tumor cells were collected, centrifuged, and resuspended in 0.9% normal saline (NS). Then, W256 tumor cells were serially diluted to a concentration of 10^2^/mL.

The rats were randomly divided into the following two groups: rats subcutaneously inoculated with tumor cells (n = 9) and control rats (n = 3). In the experimental group, rats were inoculated with 10 W256 cells in 100 µL of NS into the right flank of the rats. The control rats were subcutaneously inoculated with an equal volume of NS. All rats were anesthetized with sodium pentobarbital solution (4 mg/kg) before inoculation.

### Urine collection

Before urine collection, all rats were accommodated in metabolic cages for 2-3 days. Urine samples were collected from rats subcutaneously inoculated with tumor cells (n = 9) on days 0, 13 and 21. All rats were placed in metabolic cages individually for 12 h to collect urine without any treatment. After collection, urine samples were stored immediately at −80°C.

### Extraction and digestion of urinary proteins

Urine samples (n = 27) were centrifuged at 12,000 × g for 30 min at 4°C. Then, the supernatants were precipitated with three volumes of ethanol at −20°C overnight. The pellets were dissolved sufficiently with lysis buffer (8 mol/L urea, 2 mol/L thiourea, 50 mmol/L Tris, and 25 mmol/L DTT). After centrifugation at 4°C and 12,000 × g for 30 min, the protein samples were measured by using the Bradford assay. A total of 100 µg of each protein sample was digested with trypsin (Trypsin Gold, Mass Spec Grade, Promega, Fitchburg, Wisconsin, USA) by using filter-aided sample preparation (FASP) methods^[12]^. These digested peptides were desalted using Oasis HLB cartridges (Waters, Milford, MA) and then dried by vacuum evaporation (Thermo Fisher Scientific, Bremen, Germany).

### LC-MS/MS analysis

Digested peptides (n = 27) were dissolved in 0.1% formic acid to a concentration of 0.5 µg/µL. For analysis, 1 µg peptide from each sample was loaded into a trap column (75 µm × 2 cm, 3 µm, C18, 100 Å) at a flow rate of 0.25 µL/min and then separated with a reversed-phase analytical column (75 µm × 250 mm, 2 µm, C18, 100 Å). Peptides were eluted with a gradient extending from 4%–35% buffer B (0.1% formic acid in 80% acetonitrile) for 90 min and then analyzed with an Orbitrap Fusion Lumos Tribrid Mass Spectrometer (Thermo Fisher Scientific, Waltham, MA). The MS data were acquired using the following parameters: i) data-dependent MS/MS scans per full scan were at the top-speed mode; ii) MS scans had a resolution of 120,000, and MS/MS scans had a resolution of 30,000 in Orbitrap; iii) HCD collision energy was set to 30%; iv) dynamic exclusion was set to 30 s; v) the charge-state screening was set to +2 to +7; and vi) maximum injection time was 45 ms. Each peptide sample was analyzed twice.

### Label-free quantification

Raw data files (n=54) were searched using Mascot software (version 2.5.1, Matrix Science, London, UK) against the Swiss-Prot rat database (released in February 2017, containing 7,992 sequences). The parent ion tolerance was set to 10 ppm, and the fragment ion mass tolerance was set to 0.02 Da. The carbamidomethylation of cysteine was set as a fixed modification, and the oxidation of methionine was considered a variable modification. Two missed trypsin cleavage sites were allowed, and the specificity of trypsin digestion was set for cleavage after lysine or arginine. Dat files (n=54) were exported from Mascot software and then processed using Scaffold software (version 4.7.5, Proteome Software Inc., Portland, OR). The parameters were set as follows: both peptide and protein identifications were accepted at a false discovery rate (FDR) of less than 1.0% and proteins were identified with at least two unique peptides. Different samples were compared after normalization with the total spectra. Protein abundances at different time points were compared with spectral counting, according to previously described procedures^[13, 14]^.

### Statistical analysis

Average normalized spectral counts of each sample were used for the following statistical analysis. The levels of proteins identified on days 13 and 21 were compared with their levels on day 0. Differential proteins were selected with the following criteria: unique peptides ≥ 2; fold change ≥1.5 or ≤ 0.67; average spectral count in the high-abundance group ≥ 3; comparison between two groups were conducted using two-sided, unpaired t-test; and *P*-values of group differences were adjusted by the Benjamini and Hochberg method^[15]^. Group differences resulting in adjusted *P*-values < 0.05 were considered statistically significant. All results are expressed as the mean ± standard deviation.

### Functional enrichment analysis

Differential proteins on days 13 and 21 were analyzed by Gene Ontology (GO) based on the biological process, cellular component and molecular function categories using Database for Annotation, Visualization and Integrated Discovery (DAVID)^[16]^. The biological pathway enrichment at two time points was analyzed with IPA software (Ingenuity Systems, Mountain View, CA, USA).

## Results and discussion

### Characterization of rats subcutaneously inoculated with tumor cells

A total of 12 male Wistar rats (150 ± 20 g) were randomly divided into the following two groups: a control group (n=3) and a group of rats subcutaneously inoculated with W256 tumor cells (n=9). The body weight of these 12 rats was recorded every 3-5 days, and daily behavior changes of the two groups were observed. The body weight of the group of rats subcutaneously inoculated with W256 tumor cells was slightly lower than that of the rats in the control group, but there were no significant differences until day 41 (Figure 2). For the behavioral observations, the rats in the control group had normal daily activities with shiny hair. There were no significant differences in daily behavior between these two groups.

**Figure 2.**
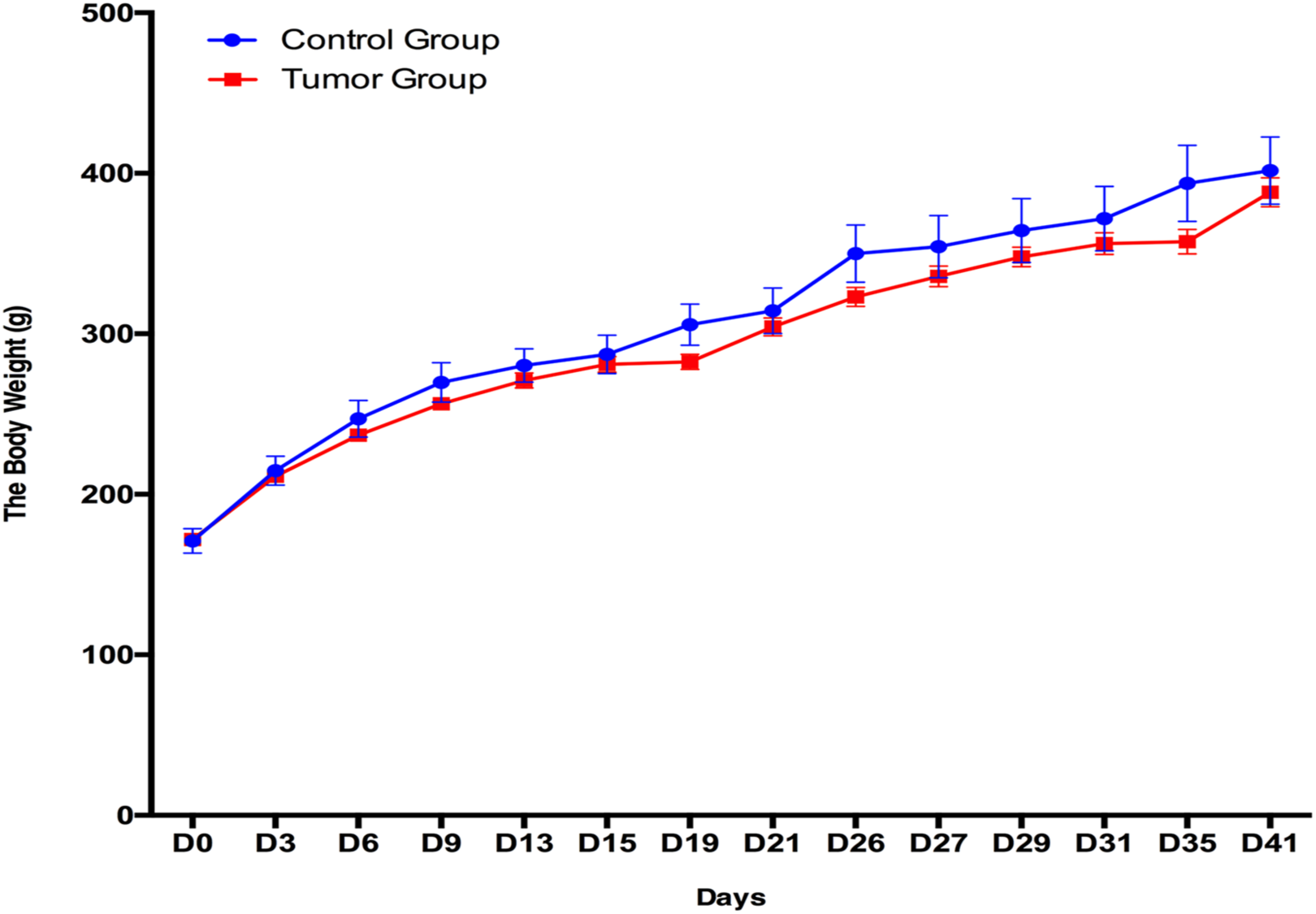
Body weight changes between the rats subcutaneously inoculated with tumor cells and the control rats.

### Urine proteome changes

Twenty-seven urine samples at three time points (days 0, 13, and 21) were used for label-free LC-MS/MS quantitation. A total of 824 urinary proteins with at least 2 unique peptides were identified with < 1% FDR at the protein level (Table S1). All identified urinary proteins were performed with unsupervised clustering analysis. Samples at three time points were almost clustered together (Figure 3A), indicating the good consistency between these nine rats. By using screening criteria, 34 and 59 differential proteins were identified on days 13 and 21, respectively. The overlap of these differential proteins is shown by a Venn diagram in Figure 3B. Details are presented in Table 1.

**Table 1.** Differentially proteins identified on day 13 and day 21.

| UniProt ID | Human ortholog | Description | Trends | ANOVA P-value | Average fold change |  |
| --- | --- | --- | --- | --- | --- | --- |
|  |  |  |  |  | Day 13 | Day 21 |
| P97603 | Q92859 | Neogenin (fragment) | ↓ | < 0.00010 | 0.43 | 0.21 |
| O70535 | P42702 | Leukemia inhibitory factor receptor | ↓ | < 0.00010 | 0.29 | 0.35 |
| Q9ESS6 | P50895 | Basal cell adhesion molecule | ↓ | < 0.00010 | 0.56 | 0.40 |
| P02761 | NO | Cluster of Major urinary protein | ↑ | < 0.00010 | 4.70 | 7.59 |
| P20611 | P11117 | Lysosomal acid phosphatase | ↓ | < 0.00010 | 0.56 | 0.46 |
| Q9JI85 | P80303 | Nucleobindin-2 | ↑ | 0.00018 | 2.74 | 3.69 |
| P07897 | P16112 | Aggrecan core protein | ↓ | < 0.00010 | 0.23 | 0.30 |
| Q63416 | Q06033 | Inter-alpha-trypsin inhibitor heavy chain H3 | ↓ | < 0.00010 | 0.32 | 0.23 |
| P06760 | P08236 | Beta-glucuronidase | ↑ | 0.0073 | 3.56 | 4.55 |
| O35112 | Q13740 | CD166 antigen | ↓ | < 0.00010 | 0.41 | 0.33 |
| Q63556 | NO | Serine protease inhibitor A3M (fragment) | ↓ | < 0.00010 | 0.47 | 0.48 |
| P26453 | P35613 | Basigin | ↓ | 0.00094 | 0.23 | 0.21 |
| P27590 | P07911 | Uromodulin | ↑ | < 0.00010 | 2.31 | 2.65 |
| P35444 | P49747 | Cartilage oligomeric matrix protein | ↓ | < 0.00010 | 0.27 | 0.42 |
| Q9EPF2 | P43121 | Cell surface glycoprotein MUC18 | ↓ | < 0.00010 | 0.18 | 0.08 |
| P13596 | P13591 | Neural cell adhesion molecule 1 | ↓ | < 0.00010 | 0.28 | 0.36 |
| P18292 | P00734 | Prothrombin | ↓ | < 0.00010 | 0.60 | - |
| P07154 | P07711 | Cathepsin L1 | ↓ | 0.00021 | 0.40 | - |
| Q63467 | P04155 | Trefoil factor 1 | ↓ | 0.00026 | 0.50 | - |
| D3ZTE0 | P00748 | Coagulation factor XII | ↓ | < 0.00010 | 0.25 | - |
| Q5HZW5 | Q9NPF0 | CD320 antigen | ↓ | < 0.00010 | 0.45 | - |
| Q8JZQ0 | P09603 | Macrophage colony-stimulating factor 1 | ↓ | 0.00013 | 0.42 | - |
| P26644 | P02749 | Beta-2-glycoprotein 1 | ↓ | 0.00016 | 0.46 | - |
| P24268 | P07339 | Cathepsin D | ↑ | < 0.00010 | 1.56 | - |
| P04937 | P02751 | Fibronectin | ↓ | < 0.00010 | 0.61 | - |
| Q63530 | Q96BW5 | Phosphotriesterase-related protein | ↑ | 0.0004 | 3.18 | - |
| P97546 | Q9Y639 | Neuroplastin | ↓ | < 0.00010 | 0.47 | - |
| P07171 | P05937 | Cluster of Calbindin | ↓ | < 0.00010 | 0.05 | - |
| P24090 | P02765 | Alpha-2-HS-glycoprotein | ↓ | 0.00026 | 0.47 | - |
| P38918 | O95154 | Aflatoxin B1 aldehyde reductase member 3 | ↑ | 0.0028 | 3.36 | - |
| P08592 | NO | Amyloid beta A4 protein | ↓ | < 0.00010 | 0.56 | - |
| Q62930 | P02748 | Complement component C9 | ↓ | 0.0055 | 0.49 | - |
| Q91XN4 | Q13145 | BMP and activin membrane-bound inhibitor homolog | ↓ | 0.003 | 0.55 | - |
| Q9JLS4 | Q6FHJ7 | Secreted frizzled-related protein 4 | ↓ | 0.00053 | 0.52 | - |
| Q5I0D5 | Q9H008 | Phospholysine phosphohistidine inorganic pyrophosphate phosphatase | ↓ | < 0.00010 | - | 0.20 |
| P80202 | P36896 | Activin receptor type-1B | ↑ | < 0.00010 | - | 1.76 |
| Q9WUW3 | P05156 | Complement factor I | ↓ | < 0.00010 | - | 0.42 |
| P07151 | P61769 | Beta-2-microglobulin | ↑ | < 0.00010 | - | 1.65 |
| Q642A7 | Q8WW52 | Protein FAM151A | ↓ | 0.00045 | - | 0.58 |
| Q63678 | P25311 | Zinc-alpha-2-glycoprotein | ↓ | < 0.00010 | - | 0.20 |
| Q9EQV6 | O14773 | Tripeptidyl-peptidase 1 | ↓ | 0.0033 | - | 0.30 |
| Q63751 | NO | Vomeromodulin (fragment) | ↓ | 0.012 | - | 0.31 |
| P04073 | P20142 | Gastricsin | ↓ | 0.00044 | - | 0.27 |
| O88917 | NO | Adhesion G protein-coupled receptor L1 | ↓ | < 0.00010 | - | 0.42 |
| P36374 | NO | Prostatic glandular kallikrein-6 | ↑ | 0.0034 | - | 1.78 |
| Q63475 | Q92932 | Receptor-type tyrosine-protein phosphatase N2 | ↓ | < 0.00010 | - | 6.25 |
| Q01460 | Q01459 | Di-N-acetylchitobiase | ↓ | < 0.00010 | - | 0.61 |
| Q99PW3 | Q99519 | Sialidase-1 | ↓ | 0.00025 | - | 0.18 |
| P85971 | O95336 | 6-phosphogluconolactonase | ↓ | 0.0005 | - | 0.39 |
| P47820 | P12821 | Angiotensin-converting enzyme | ↓ | 0.0016 | - | 0.13 |
| Q1WIM1 | Q8NFZ8 | Cell adhesion molecule 4 | ↓ | < 0.00010 | - | 0.24 |
| P05545 | NO | Serine protease inhibitor A3K | ↓ | < 0.00010 | - | 0.61 |
| P81828 | NO | Urinary protein 2 | ↑ | < 0.00010 | - | 1.96 |
| P31211 | P08185 | Corticosteroid-binding globulin | ↓ | 0.00011 | - | 0.48 |
| P04785 | P07237 | Protein disulfide-isomerase | ↓ | < 0.00010 | - | 0.17 |
| P06866 | P00739 | Haptoglobin | ↑ | 0.00036 | - | 1.68 |
| Q562C9 | Q9BV57 | 1,2-dihydroxy-3-keto-5-methylthiopentene dioxygenase | ↓ | 0.0057 | - | 0.43 |
| Q4V885 | Q5KU26 | Collectin-12 | ↑ | 0.0011 | - | 2.28 |
| P21704 | P24855 | Deoxyribonuclease-1 | ↓ | < 0.00010 | - | 0.56 |
| P63018 | P11142 | Cluster of heat shock cognate 71 kDa protein | ↓ | 0.00045 | - | 0.56 |
| P00714 | P00709 | Alpha-lactalbumin | ↑ | < 0.00010 | - | 3.77 |
| Q6AYE5 | Q86UD1 | Out at first protein homolog | ↓ | < 0.00010 | - | 0.16 |
| Q68FQ2 | Q9BX67 | Junctional adhesion molecule C | ↓ | 0.0011 | - | 0.53 |
| O55004 | P34096 | Ribonuclease 4 | ↑ | 0.00023 | - | 2.13 |
| Q641Z7 | Q92484 | Acid sphingomyelinase-like phosphodiesterase 3a | ↓ | < 0.00010 | - | 0.18 |
| O70244 | O60494 | Cubilin | ↓ | < 0.00010 | - | 0.25 |
| Q00657 | Q6UVK1 | Chondroitin sulfate proteoglycan 4 | ↓ | < 0.00010 | - | 0.28 |
| P29598 | P00749 | Urokinase-type plasminogen activator | ↓ | < 0.00010 | - | 0.62 |
| P01026 | P01024 | Complement C3 | ↓ | 0.0071 | - | 0.42 |
| P53813 | P07225 | Vitamin K-dependent protein S | ↓ | 0.00096 | - | 0.29 |
| Q80WD1 | Q86UN3 | Reticulon-4 receptor-like 2 | ↓ | 0.017 | - | 0.04 |
| P62986 | P62987 | Ubiquitin-60S ribosomal protein L40 | ↑ | 0.0011 | - | 1.81 |
| Q9R0T4 | P12830 | Cadherin-1 | ↓ | < 0.00010 | - | 0.47 |
| P08937 | NO | Odorant-binding protein | ↑ | 0.002 | - | 1.51 |
| P05544 | NO | Serine protease inhibitor A3L | ↓ | 0.00026 | - | 0.64 |
| Q05175 | P80723 | Brain acid soluble protein 1 | ↑ | < 0.00010 | - | 3.69 |
| P61972 | P61970 | Nuclear transport factor 2 | ↓ | 0.0024 | - | 0.54 |

**Figure 3.**
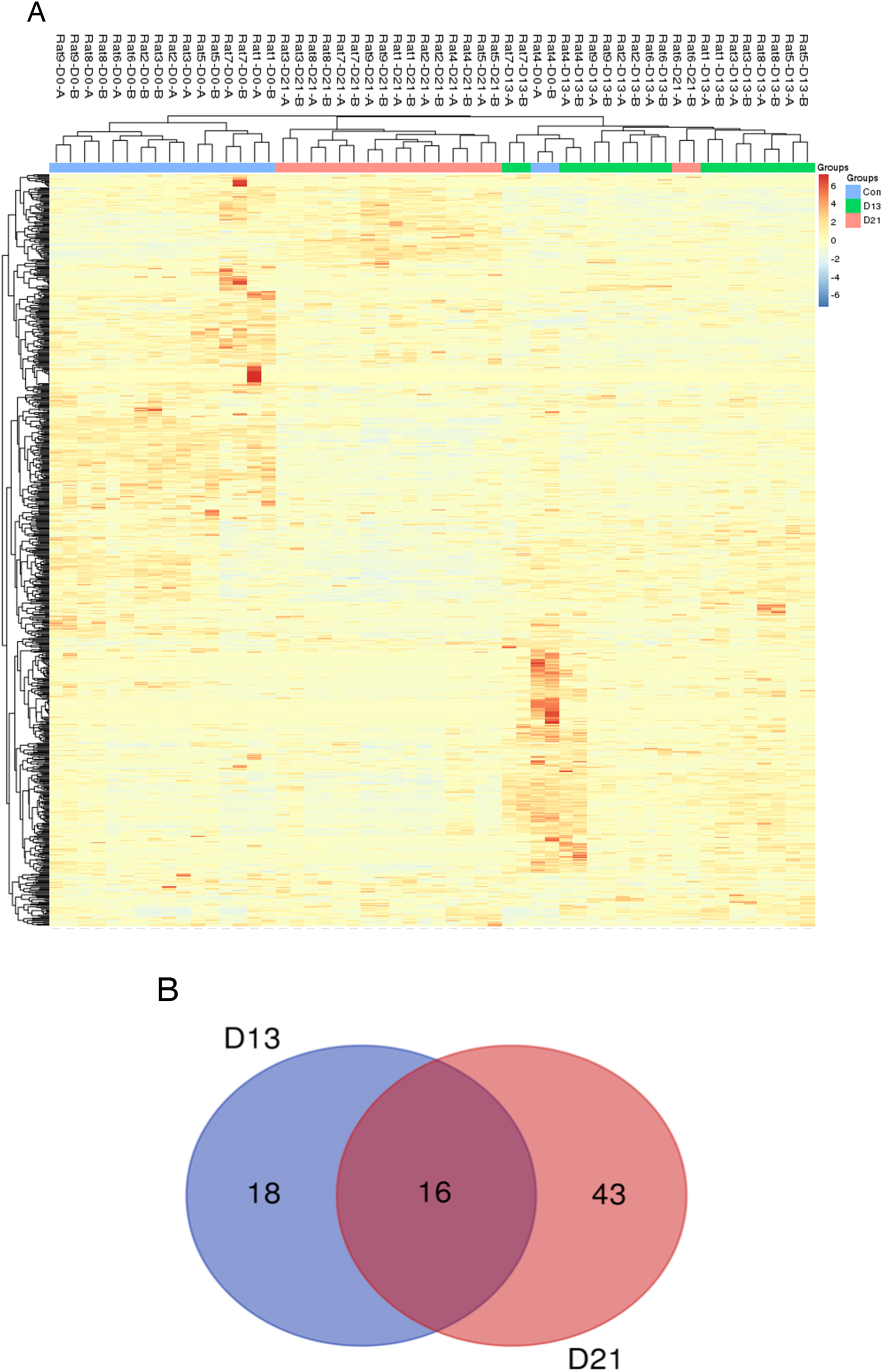
Proteomic analysis of urine samples on days 13 and 21 in rats subcutaneously inoculated with tumor cells. (A) Unsupervised cluster analysis of all proteins identified by LC-MS/MS. (B) Overlap evaluation of the differential proteins identified on days 13 and 21.

### Functional analysis

Functional enrichment analysis of differential proteins was performed by DAVID^[16]^. Differential proteins were classified into biological processes, cellular components and molecular functions. The major biological pathways of differential proteins were enriched by IPA software. A significance threshold of *P* < 0.05 was used in all these representative lists.

In the biological process, fourteen representative lists on days 13 and 21 are presented in Figure 4A. Cell adhesion, negative regulation of endopeptidase activity and organ regeneration were overrepresented both on days 13 and 21. Blood coagulation, acute-phase response, autophagic cell death, positive regulation of cell proliferation, extracellular matrix organization, response to glucose and so on were independently overrepresented on day 13. On day 21, heterophilic cell-cell adhesion via plasma membrane cell adhesion molecules, proteolysis, positive regulation of phagocytosis and angiogenesis were independently enriched. Some enriched biological processes were reported to be associated with tumor progression. For example, i) cell adhesion was usually reported to show a reduction, which correlates with tumor invasion and metastasis^[17]^. In addition, changes in cell adhesion and motility are usually considered key elements in determining the development of invasive and metastatic tumors^[18]^; ii) fibrinolytic system was also reported to have a potential pathophysiologic role in breast cancer progression^[19, 20]^; iii) autophagic cell death occurs via the activation of autophagy, which was reported to play dual roles in cancer^[21]^; iv) positive regulation of cell proliferation is a common characteristic of cancer, and inhibition of cancer cell proliferation may serve as a potential target for cancer treatment^[22]^; v) changes in extracellular matrix organization were reported with crucial roles in cancer metastasis^[23, 24]^; vi) positive regulation of blood coagulation was frequently reported in tumor progress^[25]^, even in early stages of cancer^[26]^; vii) angiogenesis was still considered as a common characteristic of tumorigenesis^[27]^, and blood vessels were considered a key target for cancer therapeutic management^[28]^; and viii) cancer cells and their surrounding microenvironment were reported to regulate pericellular proteolysis, which means the pericellular proteases contribute to cancer progression^[29]^.

**Figure 4.**
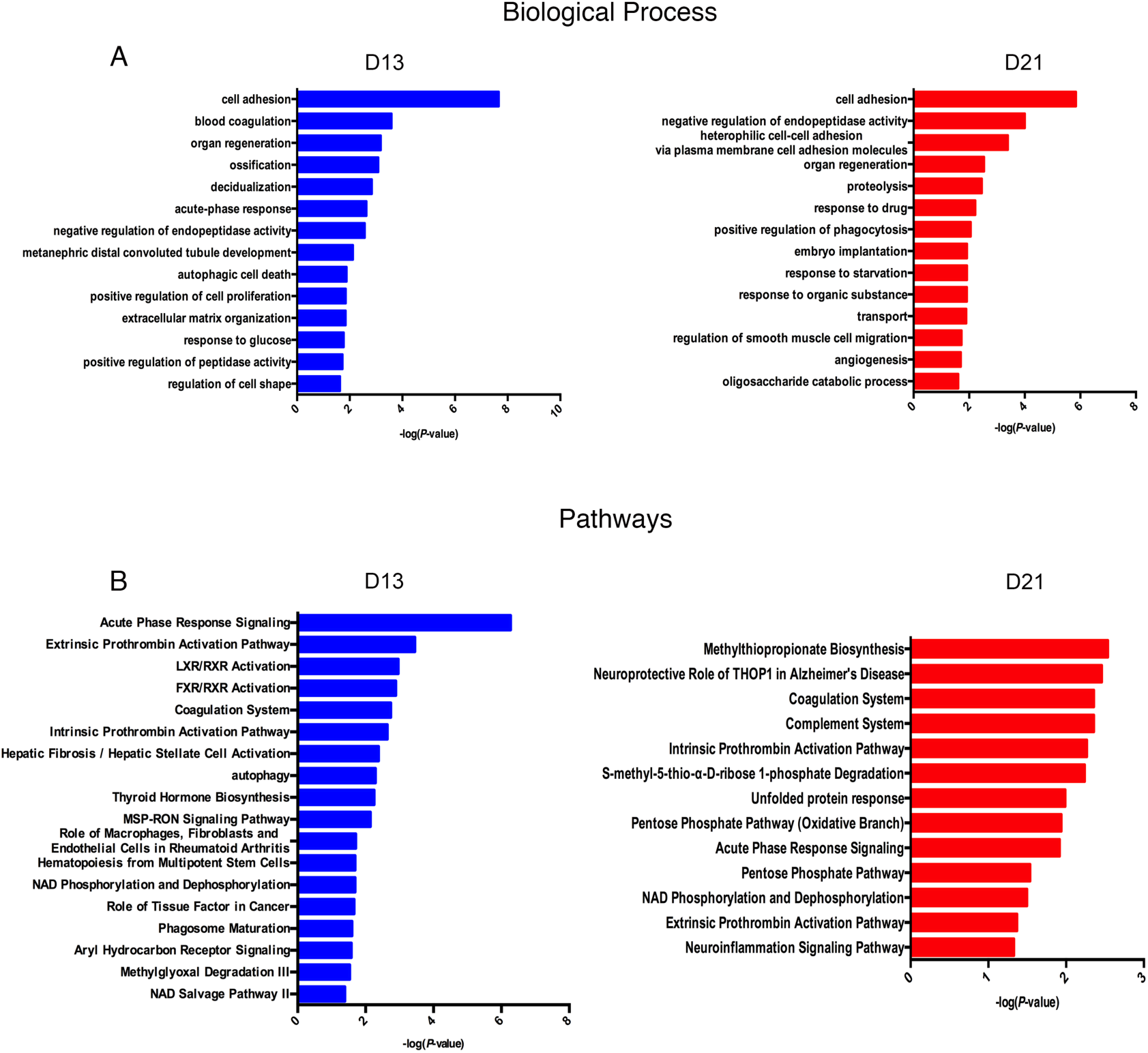
Functional analysis of differential proteins on days 13 and 21. Dynamic changes of biological processes (A) and pathways (B) at two time points were classified.

To identify the biological pathways involved with the differential urine proteins, IPA software was used for canonical pathway enrichment analysis. A total of 18 and 13 significant pathways were enriched on days 13 and 21, respectively (Figure 4B). Among these pathways, the enriched intrinsic prothrombin activation pathway, coagulation system, acute phase response signaling, extrinsic prothrombin activation pathway and NAD phosphorylation and dephosphorylation were overrepresented both on days 13 and 21. In addition, some representative pathways, such as autophagy, phagosome maturation and role of tissue factor in cancer, were independently enriched on day 13, and the complement system was enriched only on day 21. Some enriched pathways were also reported to play important roles in tumor progression. For example, i) the enriched intrinsic prothrombin activation pathway, coagulation system and extrinsic prothrombin activation pathway were all reported to participate in cancer development^[30, 31]^; ii) although in some contexts autophagy suppresses tumorigenesis, in most contexts, autophagy was reported to facilitate tumorigenesis^[32]^; iii) the MSP-RON signaling pathway was reported to play important roles in the invasive growth of different types of cancers both in vitro and in vivo^[33]^; iv) thyroid hormone biosynthesis was reported not only to play important roles in regulating normal metabolism, development, and growth but also to stimulate cancer cell proliferation^[34]^; v) tissue factor (TF) was expressed by tumor cells, which was reported to contribute to a variety of pathologic processes, such as thrombosis, metastasis, tumor growth, and tumor angiogenesis^[35, 36]^; vi) the pentose phosphate pathway (PPP) branches from glycolysis at the first committed step of glucose metabolism and was reported to play a pivotal role in helping glycolytic cancer cells to meet their anabolic demands and combat oxidative stress^[37, 38]^; and vii) the enriched complement system pathway was reported to enhance tumor growth and increased metastasis once activated in the tumor microenvironment^[39]^.

The enriched cellular components and molecular functions are presented in Figure S1. The majority of differential proteins were derived from the extracellular exosomes, extracellular space and extracellular matrix (Figure S1A). In the molecular function category, serine-type endopeptidase inhibitor activity and calcium ion binding were overrepresented both on days 13 and 21 (Figure S1B). The functional enrichment analysis indicates that even limited tumor cells exist in the body, and the urine proteome can reflect changes associated with cancer.

### Random allocation statistical analysis

To confirm that these differential proteins on days 13 and 21 were indeed due to the ten W256 tumor cells subcutaneously inoculated, we randomly allocated the data of these 27 samples (Number 1 to Number 27) into three groups. We tried three random allocations, and the numbers in these three groups are shown in Table 2. In each iteration, we used the data of group 1 as the control group. When we used the previous criteria to screen differential urinary proteins, it was found that the adjusted *P*-value value both on days 13 and 21 were all >0.05. No differential proteins were selected in these three randomly allocated trials. Details are shown in Tables S2, S3 and S4.

**Table 2.** Random allocation of the twenty-seven urine samples.

| Randomly allocated | Group 1 | Group 2 | Group 3 | Adjusted *P*-value |
| --- | --- | --- | --- | --- |
| 1 | 1, 4, 5, 12, 13,<br>14, 19, 20, 21 | 2, 3, 7, 11, 15,<br>17, 18, 22, 23 | 6, 8, 9, 10, 16,<br>24, 25, 26, 27 | NO |
| 2 | 1, 2, 3, 12, 13,<br>15, 25, 26, 27 | 4, 5, 6, 10, 11,<br>14, 19, 20, 21 | 7, 8, 9, 16, 17,<br>18, 22, 23, 24 | NO |
| 3 | 1, 7, 9, 11, 13,<br>16, 19, 21, 22 | 3, 4, 5, 10, 17,<br>18, 20, 25, 26 | 2, 6, 8, 12, 14,<br>15, 23, 24, 27 | NO |

Numbers 1-9 represent Rat 1-D0 to Rat 9-D0; Numbers 10-18 represent Rat 1-D13 to Rat 9-D13; Numbers 19-27 represent Rat 1-D21 to Rat 9-D21.

## Conclusion

In this study, we aimed to observe changes in the urine proteome when inoculating approximately ten tumor cells into nine rats. Our results indicated that 1) the urine proteome reflects changes associated with cancer, even with a limited number of tumor cells in the body, and that 2) the urine proteome reflects pathophysiological changes in the body with extremely high sensitivity, providing the potential for a very early screening process in clinical patients.

## Acknowledgments

This work was supported by the National Key Research and Development Program of China (2018YFC0910202 and 2016YFC1306300), the Beijing Natural Science Foundation (7172076), the Beijing Cooperative Construction Project (110651103), the Beijing Normal University (11100704), and the Peking Union Medical College Hospital (2016-2.27).

## Supporting Information

**Table S1.** Identification and quantitation details of the urine proteome identified in rats subcutaneously inoculated with tumor cells.

**Table S2.** Randomly allocation statistical analysis on the first trial.

**Table S3.** Randomly allocation statistical analysis on the second first trial.

**Table S4.** Randomly allocation statistical analysis on the third trial.

**Figure S1.** Functional analysis of differential proteins on day 13 and day 21. Dynamic changes of cellular component (A) and molecular function (B) at two time points were classified.

